# Improving Convolutional Network Interpretability with Exponential Activations

**DOI:** 10.1101/650804

**Authors:** Peter K. Koo, Matt Ploenzke

**Affiliations:** Howard Hughes Medical Institute, Har- vard University, Cambridge, MA; Department of Biostatistics, Harvard University, Boston, MA; Department of Data Sciences, Dana-Farber Cancer Institute, Boston, MA

## Abstract

Deep convolutional networks trained on regulatory genomic sequences tend to learn distributed representations of sequence motifs across many first layer filters. This makes it challenging to decipher which features are biologically meaningful. Here we introduce the exponential activation that – when applied to first layer filters – leads to more interpretable representations of motifs, both visually and quantitatively, compared to rectified linear units. We demonstrate this on synthetic DNA sequences which have ground truth with various convolutional networks, and then show that this phenomenon holds on *in vivo* DNA sequences.

## 1. Introduction

Convolutional neural networks (CNNs) applied to genomic sequence data have become increasingly popular in recent years (Alipanahi et al., 2015; Kelley et al., 2016; Zhou & Troyanskaya, 2015), demonstrating state-of-the-art accuracy on a wide variety of regulatory genomics prediction tasks, including transcription factor binding and chromatin accessibility. Their success has been attributed to the ability to learn features directly from the training data in a distributed manner (LeCun et al., 1998). These learned features are, in some cases, suggested to correspond to biologically-relevant sequence motifs, particularly in first convolutional layer filters (Alipanahi et al., 2015; Kelley et al., 2016).

An understanding of what a trained model has learned is then possible through attribution scores, which can be attained with perturbation methods (Alipanahi et al., 2015; Zhou & Troyanskaya, 2015) and saliency maps/gradient techniques (Simonyan et al., 2013; Shrikumar et al., 2017; Koo et al., 2018). However, the resultant attribution maps tend to be difficult to interpret, requiring downstream analysis to obtain more interpretable features, such as sequence motifs, by averaging clusters of attribution scores (Shrikumar et al., 2018). The factors that influence the quality of attribution scores – such as the CNN architecture, regularization, and training procedure – are not well characterized. There is no guarantee that attribution methods will reveal features that are biologically interpretable for a given CNN, even if it is capable of a high classification performance.

An alternative approach is to design CNNs such that their filters directly learn more interpretable features (Koo & Eddy, 2018; Ploenzke & Irizarry, 2018). In this manner, minimal posthoc analysis is required to obtain representations of “salient” features, such as sequence motifs. For instance, pre-convolution weight transformations that model the first layer filters as position weight matrices (PWMs) may be used to learn sequence motifs through the weights (Ploenzke & Irizarry, 2018). Another CNN design choice employs a large max-pool window size after the first layer, which obfuscates the spatial ordering of partial features, preventing deeper layers from heirarchically assembling them into whole feature representations (Koo & Eddy, 2018). Hence, the CNN’s first layer filters must learn whole features, because it only has one opportunity to do so.

One drawback to current design principles of CNNs with interpretable filters is that they tend to be limited to shallower networks. Depth of a network significantly increases its expressivity (Raghu et al., 2016), which enables it to learn a wider repertoire of features. In regulatory genomics, deeper networks have found greater success at classification performance. In practice, deeper CNNs are generally harder to train and are more susceptible to performance variations with different hyperparameter settings.

One consideration for the interpretability of a CNN’s filters that has not been thoroughly explored in genomics is the activation function. Rectified linear units (ReLUs) are the most commonly employed activations in genomics. In computer vision, neurons activated with a rectified polynomial, which has a close relationship to dense associative memories (Krotov & Hopfield, 2016), were shown to learn representations of numbers when applied to the MNIST dataset. This activation breaks common sense because it is unbounded and hence can diverge relatively quickly.

A divergent activation is intriguing from a signal processing perspective because it can force the network to regulate its weights such that the activity of a neuron does not blow up. For instance, if background signals are propagated through, then the rest of the network has to suppress this amplified noise in order to make accurate classification. We suspect that the network would instead opt for a simpler strategy of suppressing background signals prior to activation, thereby only propagating discriminatory signals. One drawback of the rectified polynomial, however, is that it is unclear how to select the order of the polynomial thus introducing another hyperparameter to tune.

Building upon these previous studies, we introduce a novel application of an exponential activation function. We perform systematic experiments on synthetic data that recapitulates a multi-class classification task to compare how activations of first layer filters affect representation learning of sequence motifs. We find that an exponential activation applied only to the first layer filters consistently learn whole motif representations, irrespective of the network’s depth and design. On the other hand, motif representations for CNNs that employ ReLU activations in the first layer predictively depend on CNN design. We then show that these results generalize to *in vivo* sequences.

## 2. Experimental overview

### Data

We analyzed a dataset from (Koo & Eddy, 2018), which consists of synthetic DNA embedded with known transcription factor (TF) motifs to recapitulate a multi-class classification task of identifying transcription factor binding motifs. Specifically, synthetic sequences, each 200 nucleotides long and composed of random DNA, were implanted with 1 to 5 known TF motifs, randomly selected with replacement from a pool of 12 motifs. This dataset makes a simplifying assumption that the only important pattern for a given binding event is the presence of a PWM-like motif in a sequence. Since we have ground truth for all of the relevant TF motifs, and also where they are embedded in each sequence, we can test the efficacy of the representations learned by a trained CNN.

### Models

We used two CNNs, namely CNN-50 and CNN-2 (Koo & Eddy, 2018), to learn “local” representations (whole motifs) and “distributed” representations (partial motifs), respectively. Both networks take as input a 1-dimensional one-hot-encoded sequence with 4 channels, one for each nt (A, C, G, T), and have a fully-connected (dense) output layer with 12 neurons that use sigmoid activations. The hidden layers for each model are:

#### 1. CNN-2

1. convolution (30 filters, size 19, stride 1) max-pooling (size 2, stride 2)
2. convolution (128 filters, size 5, stride 1, ReLU) max-pooling (size 50, stride 50)
3. fully-connected layer (512 units, ReLU)

#### 2. CNN-50

1. convolution (30 filters, size 19, stride 1) max-pooling (size 50, stride 50)
2. convolution (128 filters, size 5, stride 1, ReLU) max-pooling (size 2, stride 2)
3. fully-connected layer (512 units, ReLU)

#### 3. CNN-deep

1. convolution (30 filters, size 19, stride 1)
2. convolution (48 filters, size 9, stride 1, ReLU) max-pooling (size 3, stride 3)
3. convolution (96 filters, size 6, stride 1, ReLU) max-pooling (size 4, stride 4)
4. convolution (128 filters, size 4, stride 1, ReLU) max-pooling (size 3, stride 3)
5. fully-connected layer (512 units, ReLU)

All models incorporate batch normalization (Ioffe & Szegedy, 2015) in each hidden layer; dropout (Srivastava et al., 2014) with probabilities corresponding to layer1 0.1, layer2 0.1, layer3 0.5 for CNN-2 and CNN-50; and layer1 0.1, layer2 0.2, layer3 0.3, layer4 0.4, layer5 0.5 for DistNet; and *L*2-regularization on all parameters in the network with a strength equal to 1e-6.

### Training

We uniformly trained each model by minimizing the binary cross-entropy loss function with mini-batch stochastic gradient descent (100 sequences) for 100 epochs. We updated the parameters with Adam using default settings (Kingma & Ba, 2014). All reported performance metrics are drawn from the test set using the model parameters which yielded the lowest loss on the validation set. Each model was trained 5 times with different random initializations according to (He et al., 2015).

### Visualization of convolutional filters

To visualize first layer filters, we scanned each filter across every sequence in the test set. Sequences whose maximum activation was less than a cutoff of 50% of the maximum possible activation achievable for that filter were removed. A subsequence the size of the filter is taken about the max activation for each remaining sequence and assembled into an alignment. Subsequences that are shorter than the filter size due to their max activation being too close to the ends of the sequence were also discarded. A position frequency matrix was then created from the alignment and converted to a sequence logo.

### Quantitative motif comparison

The interpretability of each filter was assessed using the Tomtom motif comparison search tool (Gupta et al., 2007) to determine statistically significant matches to the 2016 JASPAR vertebrates database (Mathelier et al., 2015). Since the ground truth motifs are available for our synthetic dataset, we can test whether the CNNs have captured *relevant* motifs.

## 3. Results

To test the extent that activation functions influence representation learning by first layer filters, we trained various CNNs, namely CNN-2, CNN-50, and CNN-deep, on the synthetic dataset with 5 different initializations and used the average area under the precision recall curve (auPR) to compare performance and quantify the ability to learn sequence motifs using Tomtom (Gupta et al., 2007). For each network, we compared ReLU and exponential activations only on the first layer, while employing ReLU activations for the other hidden layers.

### Analyzing synthetic sequences

CNNs trained on the synthetic dataset show no significant differences in the auPR on held-out test sequences across models and across activations (Table 1). A visual comparison of the representations learned by first layer filters show that CNN-2 and CNN-deep do not learn sequence motifs well when employing ReLU activations (Fig. 1). This is expected because deeper layers are able to build hierarchical representations from partial motif features for these networks. Indeed less than 1% of the filters match ground truth motifs according to a Tomtom motif comparison search across 5 independent trials for each network. Nevertheless, about 60% of the filters of CNN-2 and CNN-deep have a statistically significant match to some motif in the JASPAR database, even though most of these matches are not relevant. As expected, larger max-pooling is required to yield interpretable filters for CNNs with ReLU activations (Koo & Eddy, 2018). Indeed 92% of CNN-50’s filters match ground truth motifs.

**Table 1.**
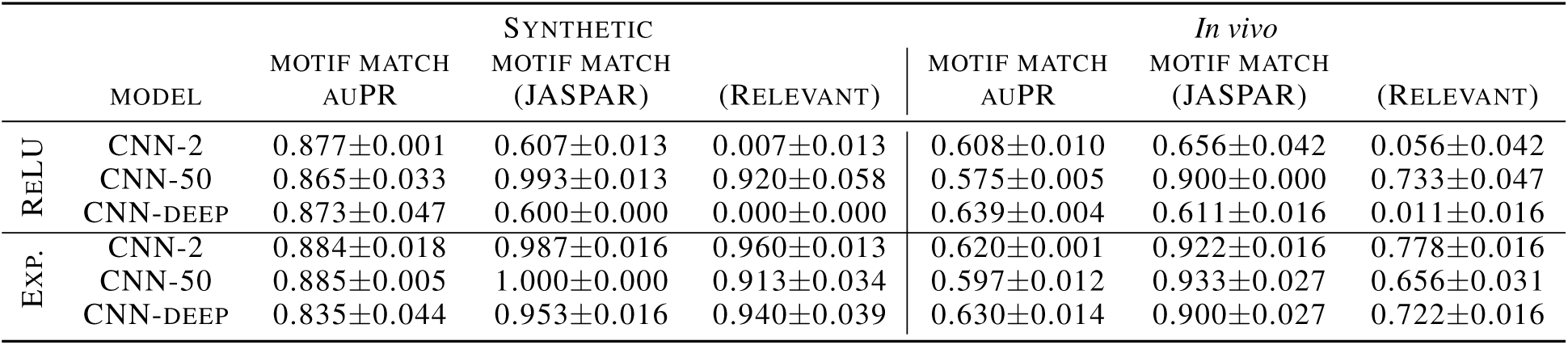
Performance comparison. This table shows the average area under the precision-recall curve (auPR) across the 12 TF classes, average percent match between the first layer filters and the entire JASPAR vertebrates database (JASPAR), and the average percent match to any ground truth TF motif (Relevant) for different CNNs. The errors represent the standard deviation across 5 independent trials.

|  | MODEL | MOTIF MATCH<br>AUPR | SYNTHETIC |  | MOTIF MATCH<br>AUPR | *In vivo* |  |
| --- | --- | --- | --- | --- | --- | --- | --- |
|  |  |  | MOTIF MATCH<br>(JASPAR) | (RELEVANT) |  | MOTIF MATCH<br>(JASPAR) | (RELEVANT) |
| ReLU | CNN-2 | 0.877±0.001 | 0.607±0.013 | 0.007±0.013 | 0.608±0.010 | 0.656±0.042 | 0.056±0.042 |
|  | CNN-50 | 0.865±0.033 | 0.993±0.013 | 0.920±0.058 | 0.575±0.005 | 0.900±0.000 | 0.733±0.047 |
|  | CNN-DEEP | 0.873±0.047 | 0.600±0.000 | 0.000±0.000 | 0.639±0.004 | 0.611±0.016 | 0.011±0.016 |
| Exp. | CNN-2 | 0.884±0.018 | 0.987±0.016 | 0.960±0.013 | 0.620±0.001 | 0.922±0.016 | 0.778±0.016 |
|  | CNN-50 | 0.885±0.005 | 1.000±0.000 | 0.913±0.034 | 0.597±0.012 | 0.933±0.027 | 0.656±0.031 |
|  | CNN-DEEP | 0.835±0.044 | 0.953±0.016 | 0.940±0.039 | 0.630±0.014 | 0.900±0.027 | 0.722±0.016 |

**Figure 1.**
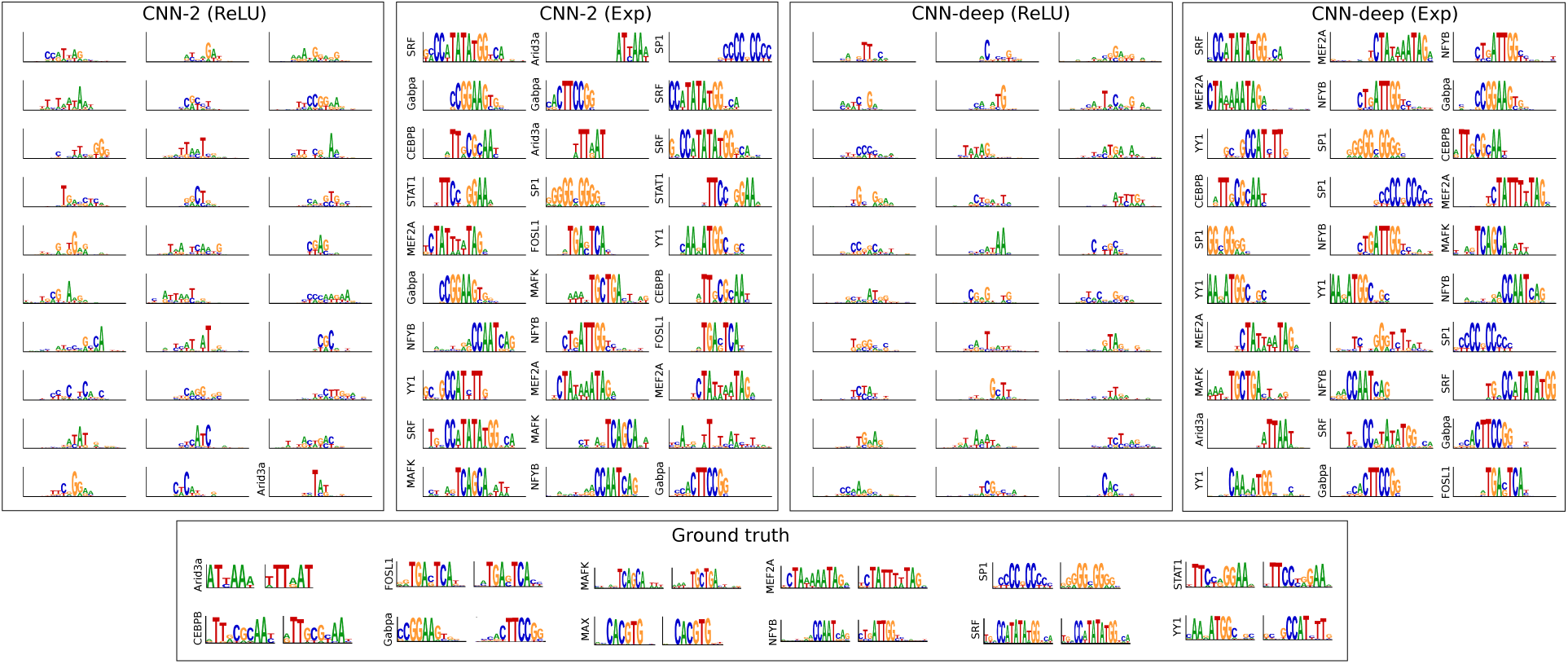
Representations learned from synthetic sequences. Sequence logos of the first convolutional layer filters are shown for (from left to right): CNN-2 with ReLU activations, CNN-2 with exponential activations, CNN-deep with ReLU activations, and CNN-deep with exponential activations. The sequence logo of the ground truth motifs and its reverse complement for each ground truth motif is shown at the bottom. The y-axis label on select filters represent a statistically significant match to a ground truth motif.

Strikingly, the convolutional filters for CNN-2 and CNN-deep, which were unable to learn motifs with ReLU activations, visually seem to capture many ground truth motifs when switching to an exponential activation (Figure 1). Quantification by Tomtom confirms that greater that 90% of the filters match ground truth motifs. This demonstrates that exponential activations provide interpretable filters for CNNs, irrespective of max-pooling size.

### Analyzing *In vivo* sequences

To test whether the same representation learning principles generalize to *in vivo* sequences, we modified the DeepSea dataset (Zhou & Troyanskaya, 2015) to include only *in vivo* sequences that have a peak called for at least one of 12 ChIP-seq experiments, each of which correspond to a TF in the synthetic dataset (see Supplemental Table S1 in (Koo & Eddy, 2018)). The truncated-DeepSea dataset is similar to the synthetic dataset, except that the input sequences now have a size of 1,000 nt in contrast to the 200 nt sequences in the synthetic dataset.

We trained each CNN on the *in vivo* dataset following the same protocol as the synthetic dataset. Similarily, a qualitative comparison of the first layer filters show that employing exponential activations consistently leads to more interpretable filters that visually matches known motifs (Fig. 2). By employing the Tomtom motif comparison search tool, we quantified the percentage of statistically significant hits between the first layer filters against the JASPAR database (see Table 1). Indeed, a higher fraction of the filters of CNNs that employ exponential activations have a statistically significant match to known motifs. On the other hand, CNNs that employ ReLU activations are more sensitive to their network design with CNN-50 being the only network that learns motifs well, yielding a percent match of 90%. We note that the performance drop for *in vivo* sequences is expected as they are more complicated, *i.e.* many filters find a GATA motif. We envision that adding more filters in the first layer can help address some of this discrepancy.

**Figure 2.**
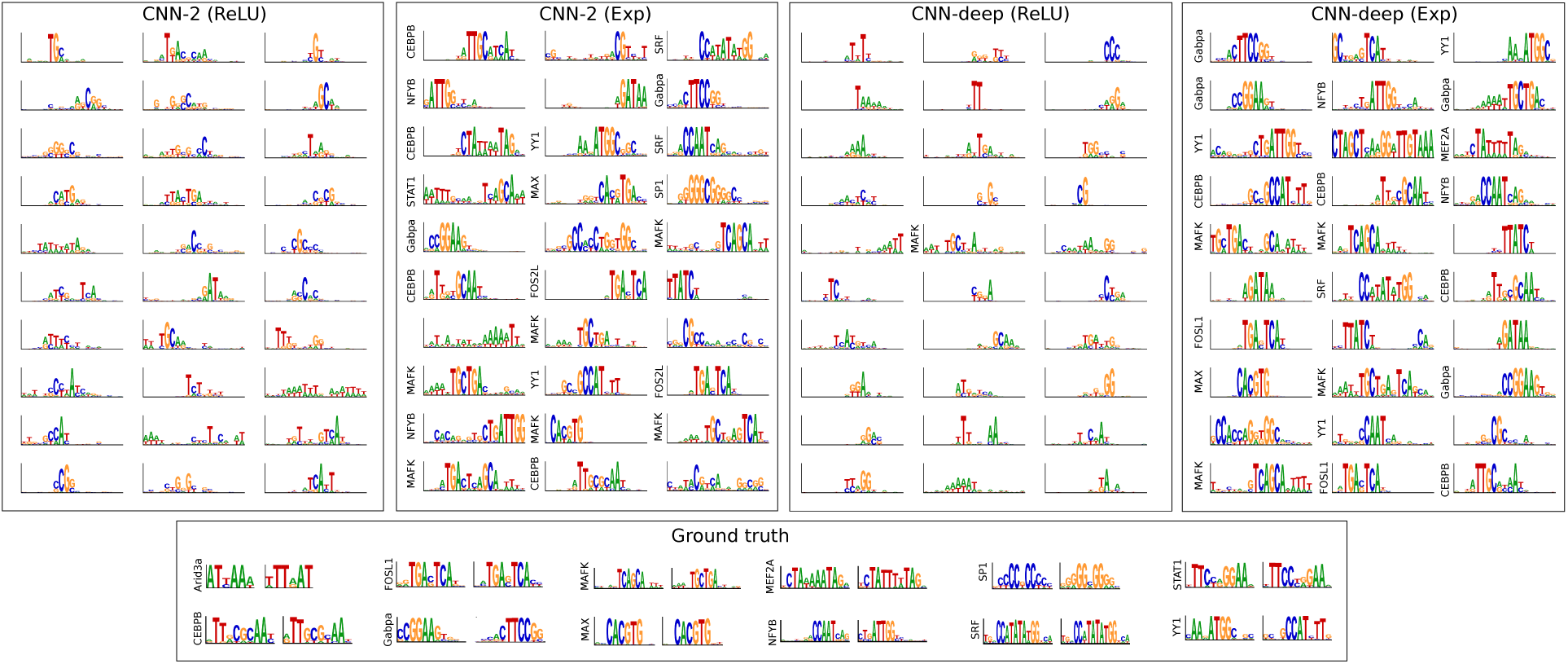
Representations learning for *in vivo* sequences. Sequence logos for first convolutional layer filters are shown for (from left to right): CNN-2 with ReLU activations, CNN-2 with exponential activations, CNN-deep with ReLU activations, and CNN-deep with exponential activations. The sequence logo of the ground truth motifs and its reverse complement for each transcription factor is shown at the bottom. The y-axis label on select filters represent a statistically significant match to a ground truth motif.

## 4. Conclusion

A major goal is to interpret learned representations of CNNs so that we can gain insights into the underlying biology. Deep CNNs, however, tend to learn distributed representations of sequence motifs that are not necessarily human interpretable. Although attribution methods can identify features that lead to decision making, their scores tend to be noisy and difficult to interpret. We show that an exponential activation is a powerful approach to encourage first layer filters to learn sequence motifs. We believe that if applied to deeper layers, it could also improve interpretability in deeper layers to potentially capture motif-motif interactions. Moving forward, one promising avenue is to combine attribution methods with CNNs that employ exponential activations so that noisy attribution scores can be aided with the interpretable first layer filters.

